# Comparative genomic analysis of *Latilactobacillus sakei* strains provides new insights into their association with different niche adaptations

**DOI:** 10.1101/2024.07.15.603503

**Authors:** Kohei Ito, Yutaro Ito

## Abstract

*Latilactobacillus sakei*, a lactic acid bacterium in diverse environments such as fermented foods, meat, and the human gastrointestinal tract, exhibits significant genetic diversity and niche-specific adaptations. This study conducts a comprehensive comparative genomic analysis of 30 complete *L. sakei* genomes to uncover the genetic mechanisms underlying these adaptations. Phylogenetic analysis divided the species into three distinct clades that did not correlate with the source of isolation and did not suggest any niche-specific evolutionary direction. The pan-genome analysis revealed a substantial core genome alongside a diverse genetic repertoire, indicating both high genetic conservation and adaptability. Predicted growth rates based on codon use bias analysis suggest that *L. sakei* strains have an overall faster growth rate and may be able to efficiently dominant in competitive environments. Plasmid analysis revealed a variety of plasmids carrying genes essential for carbohydrate metabolism, enhancing *L. sakei’*s ability to thrive in various fermentation substrates. It was also found that the number of genes belonging to the GH1 family among sugar metabolism-related genes present on chromosomes and plasmids varies between strains, and that AA1, which is involved in alcohol oxidation, has been acquired from plasmids. BLAST analysis revealed that some strains have environmental adaptation gene clusters of cell surface polysaccharides that may mediate attachment to food and mucosa. These findings not only underscore the genetic and functional diversity of *L. sakei* but also highlight its potential as a potent starter culture in fermentation and as a probiotic. The knowledge gleaned from this study lays a solid foundation for future research aimed at harnessing the genetic traits of *L. sakei* strains for industrial and health-related applications.

## Introduction

*L. sakei* has been isolated from various environments including fermented foods. Specifically, *L. sakei* has been found in meat [1], Japanese *sake* [2,3] and fermented vegetables [4]. It has also been found in human feces [5]. *L. sakei* produces lactic acid, which acidifies food products and inhibits of undesirable autochthonous microbes, allowing it to dominate in a variety of fermented foods [6]. Consequently, certain strains of *L. sakei* are used as starter cultures for the fermentation process [7,8].

*L. sakei* possesses several beneficial functions, including the production of bacteriocins that inhibit food-borne pathogenic bacteria such as *Listeria monocytogenes, Enterococcus faecalis*, and *Staphylococcus aureus* [9]. Moreover, *L. sakei* is gaining attention for its probiotic functions, including regulation of the gut environment, prevention of inflammation, and reduction of obesity [10,11].

Understanding the phenotype and adaptation mechanisms of *L. sakei* strains is crucial. Previous studies have demonstrated that environmental conditions significantly influence the genetic characteristics and evolutionary direction of bacteria. It is likely that *L. sakei* exhibits different adaptations for each fermented food. Indeed, previous studies have identified three distinct phylogenetic lineages of *L. sakei* using multilocus sequence typing (MLST), influenced by their different habitats [12]. Genes that are key for survival in meat products have been conserved in *L. sakei* isolated from these environments [13]. Large-scale comparative genomic analyses using whole genomes are necessary to further investigate the mechanisms of *L*.*sakei* adaptation to different environmental niches.

Comparative genomic analysis is a crucial method to reveal the genetic diversity and adaptation of *L. sakei* to different niches. Some studies have highlighted the role of codon usage bias in bacterial genomes as an indicator of environmental niche adaptation [14,15]. The previous study [16] have carried out comparative genomic analysis of *L. sakei* strains, but have only focused on the chromosomal genome and have not considered the effect of plasmids on environmental adaptation. In this study, we analyzed the pan-genome, phenotype prediction, CRISPR-Cas systems, and plasmids of 30 publicly available complete genomes of *L. sakei* from the National Center for Biotechnology Information (NCBI) using bioinformatics approaches.

## Materials and Methods

### Genome sequencing, assembly and annotation

For comparative analysis, RefSeq data for 30 publicly available complete genomes of *L. sakei* and some bacteria used for comparison of codon bias analysis were downloaded from the NCBI FTP site (https://www.ncbi.nlm.nih.gov/, accessed on 30 April 2023). In order to identify genomic features or plasmid detactions with high accuracy, only complete genomes were used for comparative analysis in this study. The final data set consisted of 30 *L. sakei* strains as shown in Table 1. Protein-coding DNA sequences (CDSs) were predicted and functional annotations (gene and product names) were assigned using Bakta (v.1.7.0) [17]. CRISPR arrays were detected using CRISPRCasFinder (version 4.2.30) [18] with the default parameters. CRISPR arrays with high evidence levels were considered as highly likely candidates, so we restricted the analysis to CRISPRs with evidence level ≥4 and containing at least one Cas protein. Functional categorization of genes was performed using annotation against the Cluster of Orthologous Groups of proteins (COGs) database based on BLAST [19].

### Codon usage analysis

To correlate strains with general lifestyle adaptation, we used the R package gRodon2 (version 2.3.0)[20] which uses codon usage bias to estimate growth rates. From a codon usage bias perspective, gRodon identifies the optimisation of highly expressed genes, which serves as a robust indicator of selection for growth rates.

### Pan-genome analysis

Pan-genomes were constructed using Panaroo (version 1.3.3) [21] in “sensitive” mode with the initial clustering stage at 98% length and the family sequence identity threshold at 70%. Core genes were defined as genes present in ≥95% of the *L. sakei* strains. To investigate whether the branch lengths in the phylogeny are associated with gene gain and loss, Panstripe R packages using a generalized linear model (GLM) was performed [22]. *P* values <0.05 were considered statistically significant. To infer their phylogenetic relationships, the identified single-copy orthologs were aligned using MAFFT (v7.526) [23], and a phylogenetic tree was reconstructed. The phylogenetic tree was plotted using UniPro UGENE (version 50.0) [24].

### Plasmid identification

MOB-suite (v 3.1.2) [25] was used for the typing and reconstruction of plasmid sequences from *L. sakei* strains. Plasmids are mobile genetic elements (MGEs), that allow bacteria to rapidly evolve and adapt to new niches through horizontally transfer of novel traits to different genetic backgrounds. The plasmid sequences output from the MOB-suite were gene-annotated in Bakta in the same way as chromosomal genomes.

### Cazymes, bacteriocin, and environmental adaptation genes annotation

CAZymes annotation was performed using the dbCAN3 annotation tool based on HMMER search [26]. In the analysis targeting the whole genome, CAZymes with a total of at least four genes per family across all species were used for the analysis. In the analysis using only plasmids, all annotated CAZymes were used. The identity of annotated genes was obtained with an e-value cutoff of 1e-10 or lower. Bacteriocins were annotated using BAGEL4 with whole genome sequencing [27]. The cell adherence, alcohol dehydrogenase, acid-tolerance, and salt-tolerance related genes were annotated using local BLAST [28–31]. The query sequences were obtained from CR936503, CP003032.1, and RBAI00000000. The heatmap was generated using matplotlib 3.8.4 and seaborn 0.13.2. [32].

## Results

### Genomic features of *L. sakei* strains

To date, 30 complete genomes of *L. sakei* strains have been isolated from different environments, such as human feces, fermented food, meat and milk (Table 1). The genome size of 30 *L. sakei* strains ranged from 1.88 Mbp (strain 23K) to 2.12 Mbp (strain WiKim0095) with an average size of 2.02 Mb. The average G+C content was 41.14 %, ranging from 40.97 % (strain CNSC001WB) to 41.31 % (strain J54).

All *L. sakei* strains were analysed using the CRISPR–Cas system, and only 3 strains were identified as CRISPRs at the Evidence level 4. Three strains had a complete CRISPR–Cas system (strains DS4, FLEC01 and J54), all of which belonged to class IIA or IIC, including Cas1, Cas2, Cas9, and Csn2 (Table S1).

To investigate more information about the replication abilities of *L. sakei* strains, we performed codon usage bias analysis using gRodon2, a CUB-based tool, to calculate the minimal doubling times (MDT) in hours (Figure 1A). The mean of the codon usage bias of each highly expressed gene relative to all other genes (CUBHE) of *L. sakei* strains ranged from 0.797 (strain LK-145) to 0.820 (strain DS4) with an average CUBHE of 0.810. MDT of *L. sakei* strains ranged from 0.803 (strain FLE01) to 1.116 (strain LK-145) with an average MDT of 1.009. MDT and CUBHE were negatively correlated in *L. sakei*. Comparison of MDT with a group of bacteria found in fermented foods in a previous study [33] showed that *L. sakei* has a MDT that is not particularly low compared to other bacteria, although the differences in MDTs between strains are small (Figure 1B).

**Figure 1.**
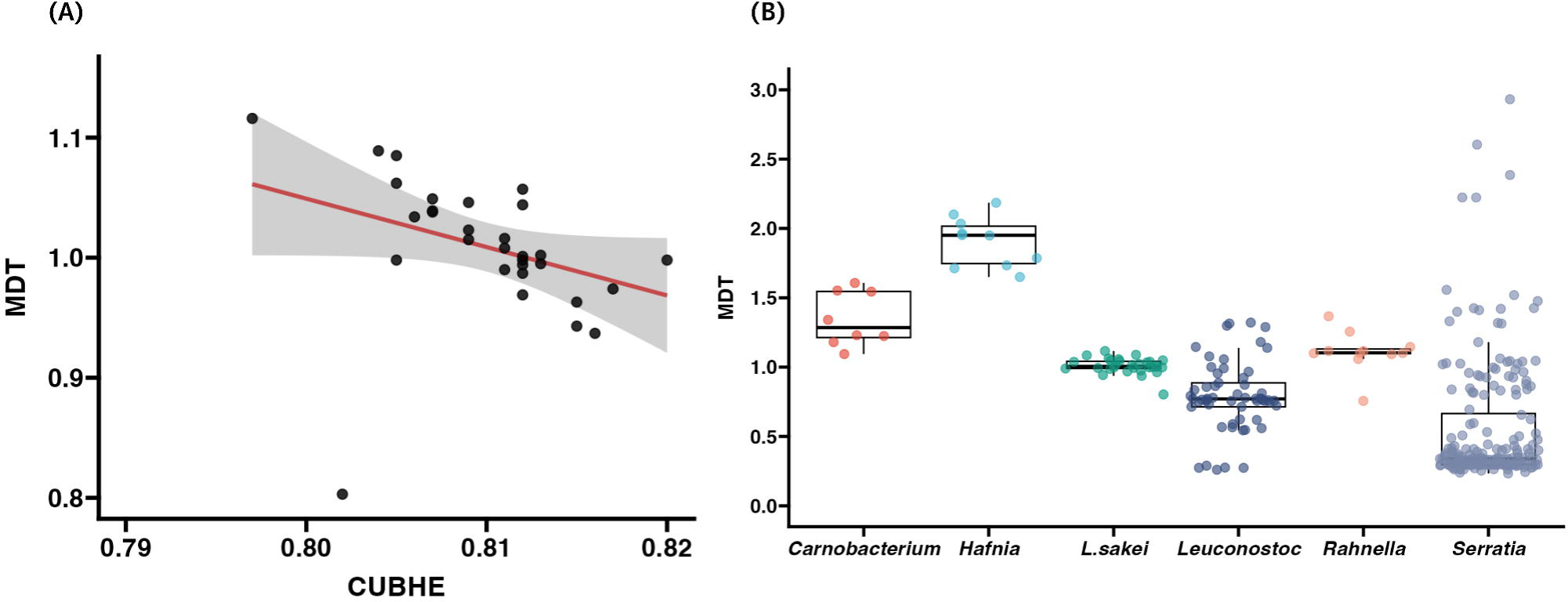
(A) Dot plot of MDT and the mean of CUBHE. (B) Boxplot of MDT between some fermented food associated bacteria.

### Pan-genome analysis

To investigate the genetic diversity among *L. sakei* strains, the number of core genes and pan-genomes and the number of strains was plotted (Figure 2A). It was shown that as the number of *L. sakei* strains increased, the number of pan-genes increased continuously and the number of core genes tended to be stable. When the 30th strain was added, the total number of genes was stable at 3,968, and the number of core genes reached 1,536. Since the results of pan-genome analysis are heavily influenced by annotation errors, we considered whether there is evidence for a temporal signal in the gene gain and loss pattern. Specifically, after fitting a panstripe model to the pan-genome, we assessed the significance of the temporal signal by calculating *p* values of the association between core branch length and gene increase/decrease. The *p* value indicates that there is a significant association between core branch lengths and gene gain/loss (*p* value > 0.005). These results show an asymptotic trend indicated that *L. sakei* has an open pan-genome.

**Figure 2.**
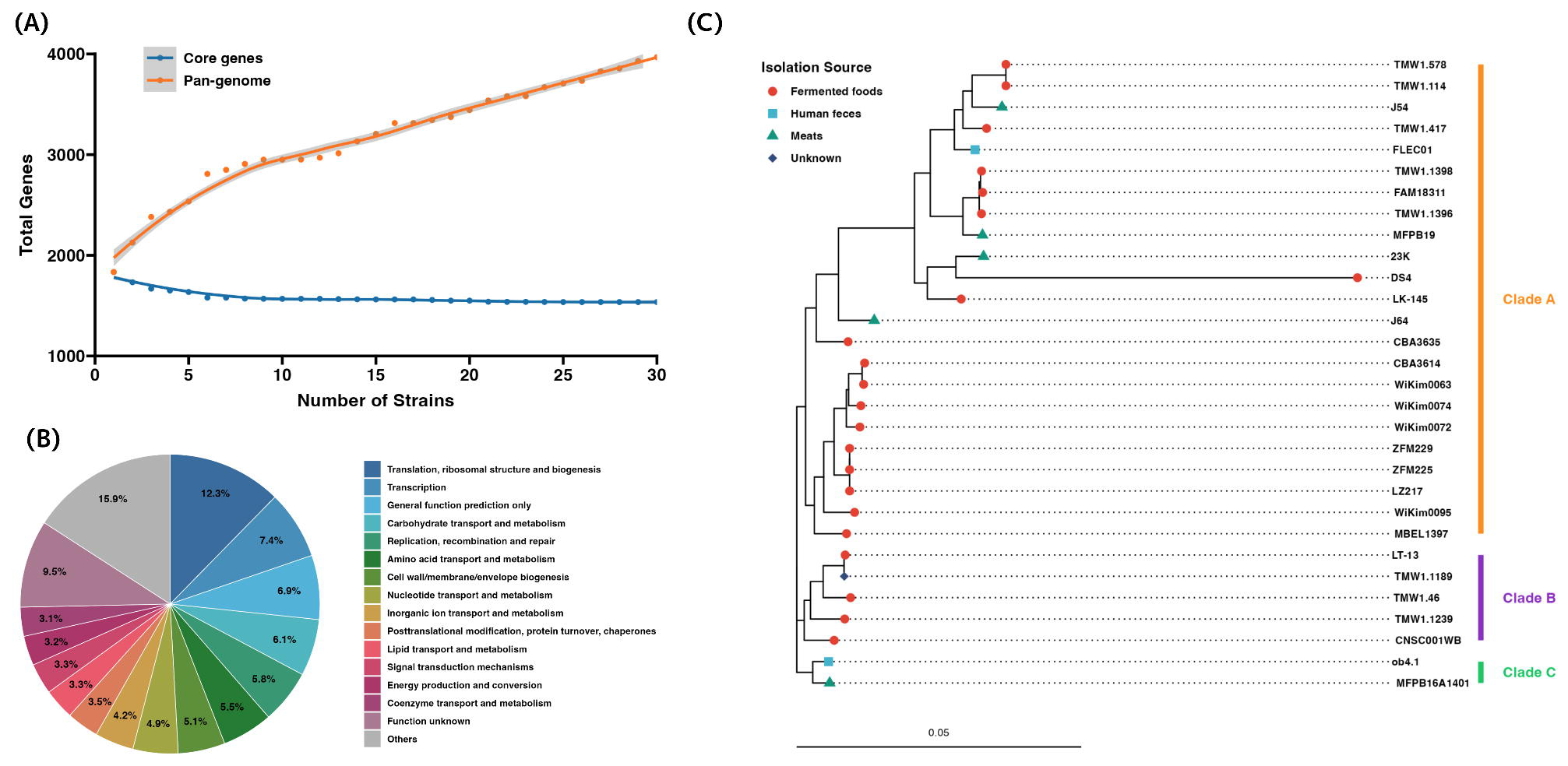
Pan-genome analysis of *L. sakei* strains. (A) Pan-genome and core gene set plots. (B) pie chart based on the database of COGs. (C) Phylogenetic tree based on the core genes set. Clade A, B and C are shown in green, blue, and yellow, respectively.

Functional analysis of the core genes of *L. sakei* strains revealed that their core genome contains more than 5.0 % of “Translation, ribosomal structure and biogenesis”, “Transcription”, “Carbohydrate transport and metabolism”,”Replication, recombination and repair”, “Amino acid transport and metabolism”, and “Cell wall/membrane/envelope biogenesis”. Among them, genes related to “Carbohydrate transport and metabolism” accounted for 6.1% of core genes, 5.68% of core genes were related to “Amino acid transport and metabolism”; however, 9.5% of core genes functions are “Function unknown” and 15.9% of core genes functions are “Others” (Figure 2B).

The phylogenetic tree was constructed based on the core genes set to explore the phylogenetic relationships of 30 *L. sakei* strains (Figure 2C). The phylogenetic tree was not divided into different clades for each source of bacterial isolates. In clade A, strains were from fermented foods and meats. Clade B were isolated from Meats and Human feces, while clade C were isolated from fermented foods and unknown. *L. sakei* from human feces (strain FLEC01 and ob4.1) were departed two different clades.

### Detection of Plasmids and their functions in *L. sakei* strains

Among 30 *L. sakei* strains, 39 plasmid sequences were detected in 22 strains (Table 2). In fact, nine strains with no plasmids detected. The G+C content of sequenced plasmids is around 34% or 44%, and genome size of the plasmids ranged from 1,526 bp to 93,254 bp as shown in Table 2. Strain LT-13 has four plasmids (primary cluster id: AB482, AA917, AA692, AH249), strain MFPB19 has three plasmids (primary cluster id: AA916, AB121, AB163) and strain FLEC01 has three plasmids (primary cluster id: AA916, AB121, AB163). Strain WiKim0095 isolated from Kimchi has a plasmid associated with *Latilactobacillus curvatus*. Strain CNSC001WB isolated from Watery radish Kimchi has *Lactiplantibacillus plantarum* 16.

A list of gene annotations found in more than 10 plasmids was generated (Table S2). 10 plasmids have some metabolic pathways. beta-phosphoglucomutase and maltose phosphorylase are involved in starch and sucrose metabolism. Aldose 1-epimerase is involved in the glycolytic system and glycogenesis.

### Genotype/Phenotype association analysis to estimate adaptation to different environments

CAZymes play a role in the synthesis of sugar complexes, oligosaccharides, and polysaccharides, aswell as the decomposition of complex carbohydrates. Some strains derived from fermented foods and from meat were each clustered into their own clade based on the number of genes in the CAZy family(Figure 3A). GT8, GH73, GT51, GH1, GT2, and GT4 were the main CAZy families in the whole genome of *L. sakei.* Some CAZymes, such as AA1, GH65, GH1, GH70, GH32, and GH91, existed on the plasmid (Figure 3B).

**Figure 3.**
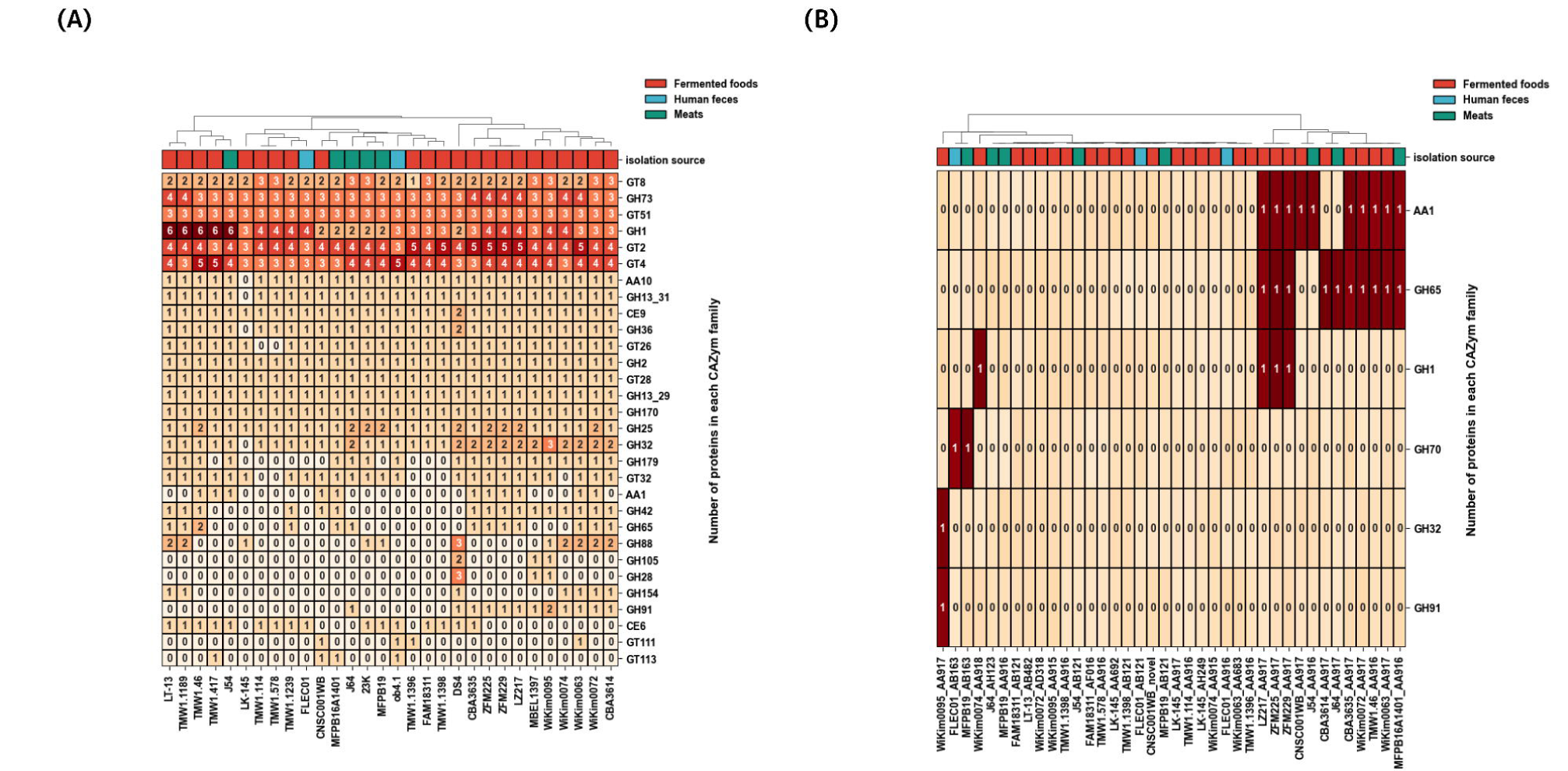
Heatmap of CAZymes. (A) The whole genomes of *L. sakei* strains from each isolation source. (B) The plasmids of *L. sakei* strains from each isolation source.

In addition, the distribution of genes related to environmental adaptation (cell adhesion, alcohol dehydrogenase, acid tolerance, and salt tolerance-related genes) revealed that these gene groups are also highly conserved regardless of the source of isolation (Figure 4). Some cell adhesion-related gene clusters, potentially involved in the production of surface polysaccharides (LSA1571 to LSA1585 and LSA1510 to LSA1513), vary in presence among different strains. 11 strains such as TMW1.114 and FAM18311 suggested the potential presence of acid-tolerance genes despite their low identity. On the other hand, it was suggested that salt tolerance and alcohol dehydrogenase related genes are present in all strains. Although there were no specific characteristics based on the isolation source, several bacteriocins were detected in each strain (Table 3). The bacteriocin detected in the most strains is Carnoicin CP52, found in LT-13, TMW 1.46, MFPB 16A1401, and TMW 1.1189. Sakacin P was detected in TMW1.114, TMW1.578, and J54. Sakacin G was possessed only by FLEC01. FAM18311 exclusively possessed lactocin S and thiopeptide. These bacteriocins were found exclusively on the chromosome, not on the plasmid.

**Figure 4.**
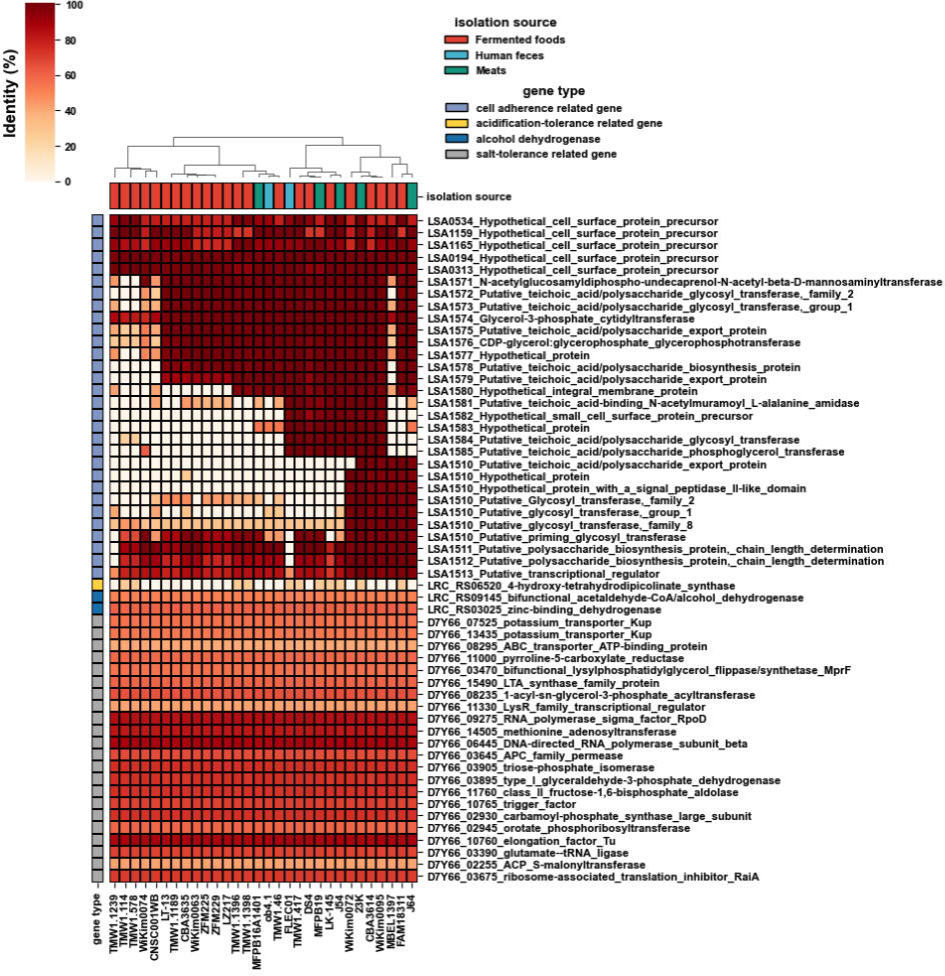
Heatmap of the environmental adaptation genes in the whole genome of *L. sakei* strains from various isolation sources.

## Discussion

The comprehensive genomic analysis of *L. sakei* strains has provided significant insights into their genetic diversity, evolutionary relationships, and adaptations to various environments. The analysis of 30 complete genomes isolated from diverse environments, including fermented food, meat, and human feces, highlights the complex and adaptive nature of *L. sakei* strains.

The variation in genome size and G+C content of *L. sakei* strains was small and consistent with previous studies [16]. Codon usage bias analysis provided insight into the replicative ability and metabolic efficiency of *L. sakei* strains (Figure 1A). The minimal doubling times (MDT) ranged from 0.803 to 1.116 hours. The doubling times of *L. sakei* strains are comparable to that of some bacteria present of final stage of the *Kimoto*-style fermentation starter [33], indicating rapid growth potential, which is advantageous for their role in fermentation processes (Figure 1B). The CUBHE values also reflected the efficiency of highly expressed genes, which are essential for the rapid adaptation and survival of *L. sakei* in competitive environments. Several studies have shown that various adventitious microbes can be detected in the fermentation process and in the production environment [33–36]. Growth rate is an important factor because lactic acid bacteria such as *L. sakei* can inhibit the growth of such adventitious microbes by multiplying rapidly and creating an acidic environment.

CRISPR-Cas systems are critical for prokaryotic adaptive immunity, countering infections by DNA and RNA viruses and foreign genes such as plasmids [37–41]. Comparative sequence analysis of *L. sakei* has identified 18 CRISPR genotypes in previous study [42]. The detection of CRISPR-Cas systems in 14 strains, with two strains possessing complete systems in *L. sakei* in this study. Active CRISPR-Cas systems are constantly adding new spacer sequences, so the number of these sequences indicates the level of activity of the system [43]. The number of spacer sequences suggests that the IIC subtype of *L. sakei* is more active and better able to resist insertion of foreign genes. The DS4 strain, which was actually the IIC subtype, had no plasmid at all. The presence of class IIA and IIC CRISPR-Cas systems suggests robust mechanisms to resist foreign genetic elements, although the absence of complete systems in most strains may confer adaptability in dynamic environments such as fermented foods.

The pan-genome analysis revealed a substantial core genome of 1,536 genes, indicating a high level of genetic conservation among the strains. The presence of 3,968 total genes underscores the extensive genetic diversity within *L. sakei*, which is likely a result of adaptation to varied environmental conditions. The core genes are primarily involved in essential metabolic processes, such as replication, transcription, translation, and various metabolic pathways (Figure 2B). This core functionality is critical for the survival and dominance of *L. sakei* in different niches. Some studies have reported that the stability and conservation of the core gene set allows bacteria to adapt to various environments [13,44]. These results suggest that *L. sakei* retains redundancy in the basic functions necessary for adaptation to diverse environments. Based on the phylogenetic tree of the core gene set, 30 *L. sakei* strains were divided into three major clades. There was no evident correlation between each branch and the isolated bacterial source (Figure 2C), consistent with the results of a previous study [16]. Interestingly, there were no significant differences in the composition of these core gene sets among strains isolated from different environments (fermented foods, meat products, human feces, etc.). This observation suggests that *L. sakei* maintains a common genetic basis that allows it to adapt to a wide range of environments.

Analysis of plasmids in *L. sakei* strains revealed significant variability, with 39 plasmid sequences detected in 22 strains. And, plasmids were also detected in J54, FLEC01 strain, which harbored the most CRISPR-Cas system genes in this study. Plasmids play an important role in horizontal gene transfer, contributing to the genetic diversity and adaptability of bacteria [45,46]. This plasmid-mediated metabolic versatility likely supports the ability of *L. sakei* to dominate in various fermentation processes and adapt to different substrates. Two *Kocuria* spp. isolated from Japanese *sake* brewing process possess plasmids with a high similarity region, indicating they share a common ancestor, and encode an identical transposase coding region, suggesting they may have migrated between isolates [35].

A list of gene annotations shared by more than 10 of all detected plasmids is shown (Table S2). It is reported that beta-phosphoglucomutase (beta-PGM) is detected in *L. plantarum*, these bacteria possess strong carbohydrate utilization capabilities, facilitated by genes such as beta-PGM, which are crucial for efficient fermentation [47]. The degradation of intracellular maltose in *Lactococcus lactis*, which is found in various lactic acid fermented foods, is known to occur through by the combined efforts of beta-phosphoglucomutase (beta-PGM) and maltose phosphorylase (MP) [48]. Aldose 1-epimerase, an enzyme that catalyzes the interconversion of the alpha- and beta-terminal isomers of hexose, is reported to be produced by Petrimonas, Prevotella and Clostridium in Chinese strongly-flavoured liquor (CSFL) produced by solid fermentation in ground pits [49]. These enzymes are important for maintaining robust fermentation activities and improving product quality.

The presence of genomes and plasmids associated with metabolic pathways, such as starch and sucrose metabolism and glycolysis, indicates their role in enhancing the metabolic capabilities of *L. sakei* (Figure 3A, B). The glycosyl hydrolases (GHs) and glycosyltransferases (GTs) have potential carbohydrate metabolic activity and play an important role in the flavor of fermented foods [50]. GH1 families including β-glucosidases and β-galactosidases, which play a crucial role in the hydrolysis of glycosidic linkages in various substrates, were found to vary in number among different strains of *L. sakei*. The GH1 family is known to be involved in lactose hydrolysis and may be present in many strains for the fermentation of milk-based products [51]. Some strains, such as Wikim0074, LZ217, ZFM225 and ZFM229, also possess one GH1 family gene in plasmid. The AA1 family, which plays an important role in the oxidation of alcohol compounds, was mainly found on plasmids. It may have been acquired from external sources to adapt to the alcohol fermentation environment.

Gene clusters involved in the production of cell surface polysaccharides, which vary among strains, may mediate attachment to both food surfaces and mucosal surfaces [28]. The genes (LSA1571 to LSA1585) predicted to be involved in the synthesis of polysaccharide-linked teichoic acid were found in strains isolated from diverse sources, indicating acquisition for various environmental adaptations. The genes (LSA1510 to LSA1513) are potentially involved in transferring polysaccharides to a surface component. These genes are primarily found in strains associated with food-related contexts, such as meat and fermented foods. As a possibility, these clusters are considered to have been acquired for colonization on food surfaces. On the other hand, many of the other adaptive genes are shared across all strains, suggesting a robust species-specific environmental adaptability.

In environmental adaptation, inhibiting the growth of other bacteria is a crucial factor. Bacteriocins are antimicrobial peptides with a proteinaceous nature, exhibiting bactericidal or bacteriostatic activity against closely related species or across genera [52]. The FAM18311 strain had Lactocin S and Thiopeptide, which exhibit activity against a broad spectrum of Gram-positive bacteria [53,54]. It is possible that they acquired it to suppress the growth of spoilage and foodborne pathogenic bacteria, thereby enhancing their environmental adaptation capabilities. Sakacin P, known as a class IIa bacteriocin produced by several *L. sakei* strains, was also present in the TMD111.4, TMD1.578, and J54 strains in this study. Sakacin P was known to exhibit activity against the food pathogen Listeria monocytogenes, thereby believed to provide an advantage in the food environment by suppressing pathogenic microorganisms [55]. As previously reported, Sakacin P is encoded on plasmids [56], indicating that bacteriocins are often acquired from plasmids as part of environmental adaptation. However, in the strains examined in this study, all bacteriocins were encoded on the chromosome. Based on these results, it was suggested that *L. sakei* can inhibit the growth of bacteria that produce various fermentations in diverse environments, implying its potential for wide-ranging applications in the food industry.

The findings from this study highlight the genetic diversity, adaptive mechanisms, and functional capabilities of *L. sakei* strains. The genomic and plasmid analyses provide valuable insights into how *L. sakei* strains have evolved to thrive in different environments, particularly in fermented foods. Understanding these adaptive traits is essential to optimize the use of *L. sakei* as a starter culture in fermentation processes and to develop strategies to harness its probiotic potential. Future research should focus on exploring the functional implications of the identified genes and plasmids in more detail to fully elucidate the mechanisms underlying the adaptability and probiotic benefits of *L. sakei* strains.

## Acknowledgements

All authors thank Morgenrot Inc. for providing the computational environment for the analysis.

---

Table 1. Genomic information of *L*.*sakei* strains.

Table 2. Plasmid genome features of *L*.*sakei* strains

Table 3. Bacteriocin in *L*.*sakei* strains

## Supplementary materials

**Table S1**. CRSIPR-Cas annotation list

**Table S2**. Plasmid annotation list

## Notes

### Competing Interest Statement

KI is a board member at BIOTA Inc., Tokyo, Japan. YI is employed by BIOTA Inc. as a part-time developer. All other authors do not have any competing interests.

### Summary of Updates

We modified the order of figures and abstract.

## Refferences

1. Hammes WP, Bantleon A, Min S. Lactic acid bacteria in meat fermentation. FEMS Microbiol Lett. 1990;87: 165–174.

2. Tsuji A, Kozawa M, Tokuda K, Enomoto T, Koyanagi T. Robust Domination of Lactobacillus sakei in Microbiota During Traditional Japanese Sake Starter Yamahai-Moto Fermentation and the Accompanying Changes in Metabolites. Curr Microbiol. 2018;75: 1498–1505.

3. Takahashi M, Morikawa K, Kita Y, Shimoda T, Akao T, Goto-Yamamoto N. Changes in Bacterial and Chemical Components and Growth Prediction for Lactobacillus sakei during Kimoto-Style Fermentation Starter Preparation in Sake Brewing: a Comprehensive Analysis. Appl Environ Microbiol. 2021;87. doi:10.1128/AEM.02546-20

4. Zhang S, Zhang Y, Wu L, Zhang L, Wang S. Characterization of microbiota of naturally fermented sauerkraut by high-throughput sequencing. Food Sci Biotechnol. 2023;32: 855–862.

5. Dal Bello F, Walter J, Hammes WP, Hertel C. Increased complexity of the species composition of lactic acid bacteria in human feces revealed by alternative incubation condition. Microb Ecol. 2003;45: 455–463.

6. Champomier-Vergès M-C, Chaillou S, Cornet M, Zagorec M. Erratum to “Lactobacillus sakei: recent developments and future prospects” [Research in Microbiology 152 (2001) 839]. Res Microbiol. 2002;153: 115–123.

7. Hammes WP, Hertel C. New developments in meat starter cultures. Meat Sci. 1998;49S1: S125–38.

8. Mani-López E, Ramírez-Corona N, López-Malo A. Latilactobacillus sakei as a starter culture to ferment pepper fruits. Food and Humanity. 2024;2: 100233.

9. Martín I, Barbosa J, Pereira SIA, Rodríguez A, Córdoba JJ, Teixeira P. Study of lactic acid bacteria isolated from traditional ripened foods and partial characterization of their bacteriocins. LWT. 2023;173: 114300.

10. Lim S, Moon JH, Shin CM, Jeong D, Kim B. Effect of Lactobacillus sakei, a Probiotic Derived from Kimchi, on Body Fat in Koreans with Obesity: A Randomized Controlled Study. Endocrinol Metab (Seoul). 2020;35: 425–434.

11. Tatsinkou LLT, Fossi BT, Sotoing GT, Mambou HMAY, Ivo PEA, Achidi EA. Prophylactic effects of probiotic bacterium Latilactobacillus sakei on haematological parameters and cytokine profile of mice infected with Plasmodium berghei ANKA during early malaria infection. Life Sci. 2023;331: 122056.

12. Chaillou S, Lucquin I, Najjari A, Zagorec M, Champomier-Vergès M-C. Population genetics of Lactobacillus sakei reveals three lineages with distinct evolutionary histories. PLoS One. 2013;8: e73253.

13. Nyquist OL, McLeod A, Brede DA, Snipen L, Aakra Å, Nes IF. Comparative genomics of Lactobacillus sakei with emphasis on strains from meat. Mol Genet Genomics. 2011;285: 297–311.

14. Botzman M, Margalit H. Variation in global codon usage bias among prokaryotic organisms is associated with their lifestyles. Genome Biol. 2011;12: R109.

15. Willenbrock H, Friis C, Juncker AS, Ussery DW. An environmental signature for 323 microbial genomes based on codon adaptation indices. Genome Biol. 2006;7: R114.

16. Chen Y, Li N, Zhao S, Zhang C, Qiao N, Duan H, et al. Integrated Phenotypic-Genotypic Analysis of from Different Niches. Foods. 2021;10. doi:10.3390/foods10081717

17. Schwengers O, Jelonek L, Dieckmann MA, Beyvers S, Blom J, Goesmann A. Bakta: rapid and standardized annotation of bacterial genomes via alignment-free sequence identification. Microb Genom. 2021;7: 000685.

18. Couvin D, Bernheim A, Toffano-Nioche C, Touchon M, Michalik J, Néron B, et al. CRISPRCasFinder, an update of CRISRFinder, includes a portable version, enhanced performance and integrates search for Cas proteins. Nucleic Acids Res. 2018;46: W246–W251.

19. Galperin MY, Makarova KS, Wolf YI, Koonin EV. Expanded microbial genome coverage and improved protein family annotation in the COG database. Nucleic Acids Res. 2015;43: D261–9.

20. Weissman JL, Hou S, Fuhrman JA. Estimating maximal microbial growth rates from cultures, metagenomes, and single cells via codon usage patterns. Proc Natl Acad Sci U S A. 2021;118: e2016810118.

21. Tonkin-Hill G, MacAlasdair N, Ruis C, Weimann A, Horesh G, Lees JA, et al. Producing polished prokaryotic pangenomes with the Panaroo pipeline. Genome Biol. 2020;21: 180.

22. Tonkin-Hill G, Gladstone RA, Pöntinen AK, Arredondo-Alonso S, Bentley SD, Corander J. Robust analysis of prokaryotic pangenome gene gain and loss rates with Panstripe. Genome Res. 2023;33: 129–140.

23. Katoh K, Standley DM. MAFFT multiple sequence alignment software version 7: improvements in performance and usability. Mol Biol Evol. 2013;30: 772–780.

24. Okonechnikov K, Golosova O, Fursov M, UGENE team. Unipro UGENE: a unified bioinformatics toolkit. Bioinformatics. 2012;28: 1166–1167.

25. Robertson J, Nash JHE. MOB-suite: software tools for clustering, reconstruction and typing of plasmids from draft assemblies. Microb Genom. 2018;4. doi:10.1099/mgen.0.000206

26. Zheng J, Ge Q, Yan Y, Zhang X, Huang L, Yin Y. dbCAN3: automated carbohydrate-active enzyme and substrate annotation. Nucleic Acids Res. 2023;51: W115–W121.

27. van Heel AJ, de Jong A, Song C, Viel JH, Kok J, Kuipers OP. BAGEL4: a user-friendly web server to thoroughly mine RiPPs and bacteriocins. Nucleic Acids Res. 2018;46: W278–W281.

28. Chaillou S, Champomier-Vergès M-C, Cornet M, Crutz-Le Coq A-M, Dudez A-M, Martin V, et al. The complete genome sequence of the meat-borne lactic acid bacterium Lactobacillus sakei 23K. Nat Biotechnol. 2005;23: 1527–1533.

29. Pang X, Li W, Yang L, Hu C, Lu J, Zhang S, et al. Whole-genome sequencing and genomic-based acid tolerance mechanisms of Lactobacillus delbrueckii subsp. bulgaricus LJJ. Research Square. 2019. doi:10.21203/rs.2.16590/v1

30. Yao W, Yang L, Shao Z, Xie L, Chen L. Identification of salt tolerance-related genes of Lactobacillus plantarum D31 and T9 strains by genomic analysis. Ann Microbiol. 2020;70: 1–14.

31. Altschul SF, Gish W, Miller W, Myers EW, Lipman DJ. Basic local alignment search tool. J Mol Biol. 1990;215: 403–410.

32. Hunter JD. Matplotlib: A 2D Graphics Environment. Comput Sci Eng. May-June 2007;9: 90–95.

33. Ito K, Niwa R, Yamagishi Y, Kobayashi K, Tsuchida Y, Hoshino G, et al. A unique case in which Kimoto-style fermentation was completed with Leuconostoc as the dominant genus without transitioning to Lactobacillus. J Biosci Bioeng. 2023;135: 451–457.

34. Ito K, Niwa R, Kobayashi K, Nakagawa T, Hoshino G, Tsuchida Y. A dark matter in sake brewing: Origin of microbes producing a Kimoto-style fermentation starter. Front Microbiol. 2023;14: 1112638.

35. Terasaki M, Kimura Y, Yamada M, Nishida H. Genomic information of Kocuria isolates from sake brewing process. AIMS Microbiol. 2021;7: 114–123.

36. Kanamoto E, Terashima K, Shiraki Y, Nishida H. Diversity of Bacillus Isolates from the Sake Brewing Process at a Sake Brewery. Microorganisms. 2021;9. doi:10.3390/microorganisms9081760

37. Barrangou R, Fremaux C, Deveau H, Richards M, Boyaval P, Moineau S, et al. CRISPR provides acquired resistance against viruses in prokaryotes. Science. 2007;315: 1709–1712.

38. Semenova E, Jore MM, Datsenko KA, Semenova A, Westra ER, Wanner B, et al. Interference by clustered regularly interspaced short palindromic repeat (CRISPR) RNA is governed by a seed sequence. Proc Natl Acad Sci U S A. 2011;108: 10098–10103.

39. Tamulaitis G, Kazlauskiene M, Manakova E, Venclovas C, Nwokeoji AO, Dickman MJ, et al. Programmable RNA shredding by the type III-A CRISPR-Cas system of Streptococcus thermophilus. Mol Cell. 2014;56: 506–517.

40. Jiang W, Samai P, Marraffini LA. Degradation of phage transcripts by CRISPR-associated RNases enables type III CRISPR-Cas immunity. Cell. 2016;164: 710–721.

41. Hatoum-Aslan A, Maniv I, Samai P, Marraffini LA. Genetic characterization of antiplasmid immunity through a type III-A CRISPR-Cas system. J Bacteriol. 2014;196: 310–317.

42. Schuster JA, Vogel RF, Ehrmann MA. Characterization and distribution of CRISPR-Cas systems in Lactobacillus sakei. Arch Microbiol. 2019;201: 337–347.

43. Tyson GW, Banfield JF. Rapidly evolving CRISPRs implicated in acquired resistance of microorganisms to viruses. Environ Microbiol. 2008;10: 200–207.

44. Lin H, Yu M, Wang X, Zhang X-H. Comparative genomic analysis reveals the evolution and environmental adaptation strategies of vibrios. BMC Genomics. 2018;19: 135.

45. Billane K, Harrison E, Cameron D, Brockhurst MA. Why do plasmids manipulate the expression of bacterial phenotypes? Philosophical Transactions of the Royal Society B: Biological Sciences. 2022;377. doi:10.1098/rstb.2020.0461

46. Heuer H, Smalla K. Plasmids foster diversification and adaptation of bacterial populations in soil. FEMS Microbiol Rev. 2012;36: 1083–1104.

47. Cui Y, Wang M, Zheng Y, Miao K, Qu X. The Carbohydrate Metabolism of Lactiplantibacillus plantarum. Int J Mol Sci. 2021;22: 13452.

48. Qian N, Stanley GA, Hahn-Hägerdal B, Rådström P. Purification and characterization of two phosphoglucomutases from Lactococcus lactis subsp. lactis and their regulation in maltose- and glucose-utilizing cells. J Bacteriol. 1994;176: 5304–5311.

49. Ren D, Liu S, Zhang S, Qin H, Han X, Mao J. Multi-omics reveals microbial roles and metabolic functions at the spatiotemporal niche in pit mud. Research Square. 2022. doi:10.21203/rs.3.rs-1251838/v1

50. Liang T, Jiang T, Liang Z, Zhang N, Dong B, Wu Q, et al. Carbohydrate-active enzyme profiles of Lactiplantibacillus plantarum strain 84-3 contribute to flavor formation in fermented dairy and vegetable products. Food Chem X. 2023;20: 101036.

51. Plaza-Vinuesa L, Sánchez-Arroyo A, Moreno FJ, de Las Rivas B, Muñoz R. Dual 6Pβ-Galactosidase/6Pβ-Glucosidase GH1 Family for Lactose Metabolism in the Probiotic Bacterium Lactiplantibacillus plantarum WCFS1. J Agric Food Chem. 2023;71: 10693–10700.

52. Gaspar C, Donders GG, Palmeira-de-Oliveira R, Queiroz JA, Tomaz C, Martinez-de-Oliveira J, et al. Bacteriocin production of the probiotic Lactobacillus acidophilus KS400. AMB Express. 2018;8: 153.

53. Cintas LM, Casaus P, Fernández MF, Hernández PE. Comparative antimicrobial activity of enterocin L50, pediocin PA-1, nisin A and lactocin S against spoilage and foodborne pathogenic bacteria. Food Microbiol. 1998;15: 289–298.

54. Darbandi A, Asadi A, Mahdizade Ari M, Ohadi E, Talebi M, Halaj Zadeh M, et al. Bacteriocins: Properties and potential use as antimicrobials. J Clin Lab Anal. 2022;36: e24093.

55. Møretrø T, Naterstad K, Wang E, Aasen IM, Chaillou S, Zagorec M, et al. Sakacin P non-producing Lactobacillus sakei strains contain homologues of the sakacin P gene cluster. Res Microbiol. 2005;156: 949–960.

56. Vaughan A, Eijsink VGH, Van Sinderen D. Functional characterization of a composite bacteriocin locus from malt isolate Lactobacillus sakei 5. Appl Environ Microbiol. 2003;69: 7194–7203.

